# Exercise restores chloride homeostasis and decreases spasticity through the BDNF-KCC2 pathway after chronic SCI

**DOI:** 10.1101/489740

**Authors:** Henrike Schulze, Samantha Choyke, Michael Klaszky, Marie-Pascale Côté

## Abstract

Activity-based therapies are routinely integrated in rehabilitation programs to induce repetitive activation of the neuromuscular system and facilitate functional recovery after spinal cord injury (SCI). Among the beneficial effects of physical therapy is a reduction of hyperreflexia and spasticity in SCI individuals, but the precise mechanism by which exercise regulates spinal networks and facilitate recovery remains elusive. Spasticity is a debilitating condition that affects ~ 75% of the SCI population and interferes with residual motor function. Current pharmacological treatments not only have serious side effects but also actively depress spinal excitability and interferes with motor recovery. Understanding how activity-based therapies contribute to decrease spasticity will help identify critical pharmacological targets and optimize rehabilitation programs.

KCC2, a neuron-specific Cl^-^ extruder, is critical to the maintenance of [Cl^-^]_i_ and its downregulation after SCI leads to a shift in chloride homeostasis that contributes to develop spasticity. We have shown in earlier studies that not only exercise promotes reflex modulation but also restores KCC2 expression in motoneurons. KCC2 is dynamically modulated by several signaling pathways, the most prevalent being BDNF-TrkB. Interestingly, activity-dependent processes triggered by exercise include an increase in the expression of BDNF in the lumbar spinal cord. However, whether the increase in KCC2 contributes to functional recovery and rely on BDNF activity have not been established. Our objective was to determine 1) whether the activity-dependent upregulation of KCC2 contributes to decrease spasticity after SCI; 2) if BDNF regulates KCC2 expression in an activity-dependent manner.

Using a model of complete SCI, we investigated this possible causal effect by intrathecally delivering VU0240551, a specific KCC2 blocker, or TrkB-IgG, a BDNF scavenger. Drugs were specifically delivered during the daily rehabilitation sessions to transiently prevent KCC2/BDNF activity. We provide evidence that the beneficial effect of exercise on functional recovery relies on a BDNF-dependent increase in KCC2 expression on motoneurons and the restoration of endogenous inhibition to a mature state. We identify, for the first time, that the increase in KCC2 activity with activity-based therapies functionally contributes to H-reflex recovery and critically depends on BDNF activity. This provides a new perspective on our understanding of how exercise impact hyperreflexia by identifying the biological basis of recovery of function. Acting directly on chloride homeostasis through BDNF to restore endogenous inhibition rather than actively depress excitability can diminish the reduction in motor output associated with the current pharmacological management of SCI and improve the outcome of rehabilitation programs.

## Introduction

Activity-based therapies promote sensorimotor functional recovery after SCI and are routinely integrated in rehabilitation programs in the clinic. Amongst its beneficial effects is a decrease in spasticity that reduces incapacitating symptoms and significantly improve the quality of life of SCI individuals (Dietz, 2001; Petropoulou *et al.*, 2007). Spasticity is a debilitating condition affecting up to 75% of SCI individuals, with most SCI individuals experiencing episodes one year after injury (Maynard *et al.*, 1990; Skold *et al.*, 1999; Holtz *et al.*, 2017). Currently available anti-spastic drugs have serious side effects including sedation, dizziness and a deep, long-lasting depression of spinal excitability that significantly reduces muscle activity and interferes with motor recovery (Dario and Tomei, 2004; Adams and Hicks, 2005; Elbasiouny *et al.*, 2010; Angeli *et al.*, 2012). There is therefore a critical need to identify new targets to diminish spasticity in SCI individuals.

Evidence-based clinical practice highlights the beneficial effect of a rehabilitation program on spastic symptoms which suggests the involvement of a potent activity-dependent mechanism. To date, the most compelling finding indicates a critical role for BDNF in promoting activity-dependent plasticity. BDNF is released in an activity-dependent manner (Lu, 2003) and its contribution to functional plasticity in the healthy and injured spinal cord has been extensively studied (reviewed in Boyce and Mendell, 2014*a*,*b*). Exercise increases BDNF serum levels in SCI individuals (Leech and Hornby, 2017). While the consequences of this activity-dependent increase remains to be determined in humans, experiments performed in animal models of chronic SCI suggest that it is associated with an enhanced response of spinal motor pools to descending drive, normalization of motoneuronal electrophysiological properties, improvement in reflex modulation and locomotor recovery, and reduction in allodynia (Hutchinson *et al.*, 2004; Beaumont *et al.*, 2008; Ying *et al.*, 2008; Côté *et al.*, 2011; Skup *et al.*, 2014).

Interestingly, BDNF acts as a regulator of the K^+^-Cl^-^ cotransporter KCC2 in various disease models including neuropathic pain and hyperalgesia (Rivera *et al.*, 2002; 2004; Coull *et al.*, 2005; Miletic and Miletic, 2008; Ferrini *et al.*, 2013). After chronic SCI, KCC2 is downregulated in spinal neurons causing an increase in intracellular chloride concentration ([Cl^-^]_i_). Consequently, GABA-mediated responses become less hyperpolarizing and lead to increased spinal reflex excitability and a significant reduction in postsynaptic inhibition (Lu *et al.*, 2008; Boulenguez *et al.*, 2010; Gackiere and Vinay, 2015). Exercise increases both BDNF and KCC2 levels in the lumbar spinal cord after a chronic SCI in animals that display less spasticity and better reflex modulation (Côté *et al.*, 2011; 2014). While it is generally accepted that activity-based therapies increase KCC2 expression in lumbar motoneurons (Côté *et al.*, 2014; Chopek *et al.*, 2015; Tashiro *et al.*, 2015), the potential contribution to functional recovery remains to be determined.

Mechanisms of activity-dependent regulation of KCC2 and subsequent shift in E_GABA_ include TrkB activation by BDNF (Aguado *et al.*, 2003; Ludwig *et al.*, 2011; Kaila *et al.*, 2014a,b). In the hippocampus, the regulation of KCC2 following neonatal status epilepticus is dependent on BDNF and accompanied by a significant increase in KCC2 expression on neuronal surface, enhancement of neuronal Cl^-^ extrusion, and hyperpolarized E_GABA_ (Khirug *et al.*, 2010; Puskarjov *et al.*, 2015). We therefore hypothesized that the beneficial effect of rehabilitation on the modulation of spinal reflexes relies on a BDNF-dependent increase in KCC2 expression in motoneurons and the restoration of endogenous inhibition.

Adult rats received a spinal transection (T12), and were exercised on motorized bikes for 4 weeks. During this exercise period, animals were treated with the specific KCC2 blocker VU0240551 or TrkB-IgG to chelate BDNF. During a terminal experiment, H-reflexes were recorded and analyzed as a measure of hyperreflexia and spasticity. Our data illustrate that preventing KCC2 activity during exercise impedes reflex recovery after SCI and that the upregulation of KCC2 expression triggered by exercise requires BDNF to restore reflex modulation. These results strongly suggest the presence of activity-dependent mechanisms involved in the regulation of chloride cotransporters in the spinal cord and demonstrate their involvement in functional recovery after SCI. This further lends support to chloride cotransporters as effective targets to improve motor recovery after SCI.

## Materials and methods

### Experimental design

In a SCI rat model of complete thoracic transection injury (T12), we investigated the role of KCC2 in motor recovery after a chronic SCI and its dependence on the activity-dependent regulation of BDNF in the lumbar spinal cord by using VU0240551, a selective inhibitor of KCC2 or chelating the endogenously released BDNF with the fusion protein TrkB-IgG. Both were delivered intrathecally during the rehabilitation program. Rats were randomly assigned to one of the following groups: 1) intact (n=11); 2) SCI sedentary control rats receiving the vehicle (SCI, n=14); 3) SCI receiving the KCC2 blocker VU0240551 (SCI+VU0240551, n=8); 4) SCI receiving the BDNF scavenger TrkB-IgG (SCI+TrkB-IgG, n=9), 5) SCI bike-trained receiving the vehicle (SCI+Ex, n=14); 6) SCI bike-trained receiving VU0240551 (SCI+Ex+VU0240551, n=14); 7) SCI-biked trained receiving TrkB-IgG (SCI+Ex+TrkB-IgG, n=11). Hyperreflexia was assessed 4 weeks after SCI and the spinal cord tissue harvested for immunohistochemistry or western blot analysis.

All procedures complied with ARRIVE, were conducted in compliance with the guidelines of the National Institutes of Health for the care and use of laboratory animals and approved by Drexel University Institutional Animal Care and Use Committee.

### Surgical procedures and postoperative care

Adult female Sprague-Dawley rats (n=106, 225-250g, Charles Rivers Laboratories) were used in this study and the timeline for procedures illustrated in Fig. 1A. All rats were housed by pairs in cages in a 12h light-dark cycle and controlled room temperature with *ad libitum* access to food. All animals but the intact group underwent a complete spinal cord transection at T12 under aseptic conditions as described previously (Côté *et al.*, 2014). Briefly, rats were anaesthetized with isoflurane (1-4%) and their back shaved, cleansed and disinfected. Body temperature was maintained ~37°C throughout the surgical procedure and post-surgical recovery. A laminectomy (T10-11) was performed, the dura slit open and a 2mm cavity created by aspiration. A second laminectomy was also performed (L2) and the proximal end of an intrathecal catheter (Alzet^®^, Durect Corporation) threaded rostrally in the subdural space over a ~10mm distance. The catheter was secured and glued onto a dry clean transverse process and the distal end coupled to a programmable microinfusion pump (iPrecio^©^, Durect Corporation) (Tan *et al.*, 2011). The pump reservoir was filled with saline and the drug delivery program activated. Back muscles were sutured in layers leaving the distal end of the catheter exiting between 2 sutures. The pump was positioned subcutaneously just below the shoulder blades and sutured to back muscles. The remaining portion of tubing was coiled for stress relief and the skin closed with staples. The completeness of the lesion was recognized by the retraction of the rostral and caudal portions of the spinal cord and inspecting the ventral floor of the spinal canal during the surgery as well as confirmed post-mortem. The animals were given saline (5mL/day s.c. for 3 days) to avoid dehydration, prophylactic cefazolin (160mg/kg/day s.c. for 7 days) and slow release buprenorphine (0.05mg/kg, s.c.) as an analgesic for pain control. Bladders were expressed manually twice a day until the end of the study.

**Figure 1.**
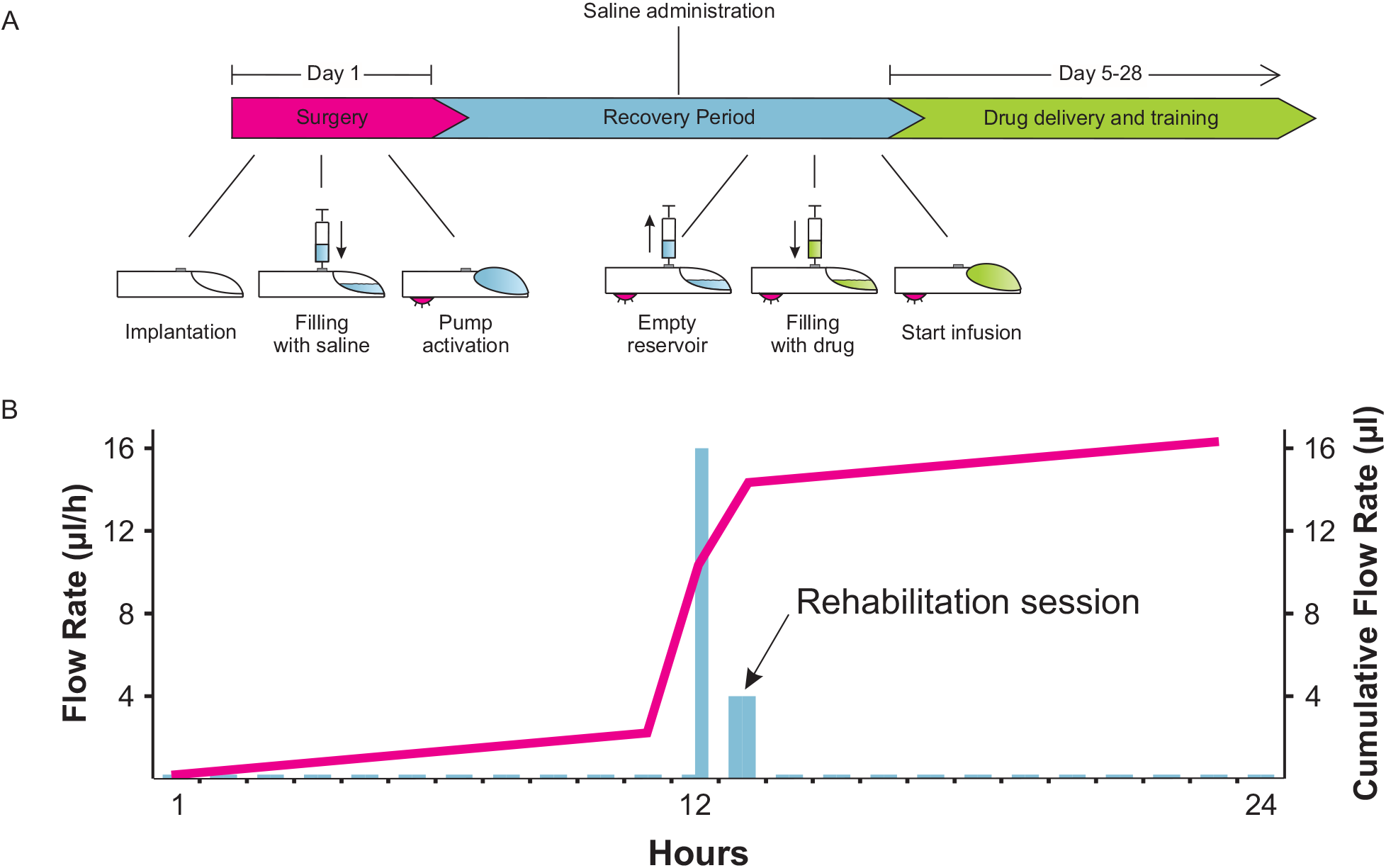
Experimental timeline and programmable infusion pump delivery. **A)** The surgical procedure took place on day 1. During the procedure, a complete spinal cord injury was performed at the thoracic level (T12). An intrathecal catheter was also implanted subdurally and fitted to a programmable microinfusion pump (iPrecio©) to deliver drugs to the lumbar enlargement of the spinal cord. During the post-SCI recovery period (~4 days), saline was continuously delivered at a steady flow rate (3μl/hour) to prevent blockage of the tip of the cannula. The iPrecio^©^ Management system calculates the time at which the reservoir needs to be emptied from saline and re-filled with drugs so as to reach the tip of the cannula 30 minutes prior to the first rehabilitation session. The saline remaining in the reservoir was then replaced through the subcutaneous port with the vehicle (10% DMSO in saline) for SCI controls, VU0240551 (30 μM) or TrkB-IgG (10 μg/ml in PBS). **B)** The pumps were programmed on a variable 24h delivery schedule. Delivery started 30 minutes before the exercise session at a flow rate of 16 μl/h for 30 minutes followed by a 4 μl/h maintenance dose during the 60 minutes training session. Drug delivery was tapered off during the remaining 22.5 hours to 0.2 μl/h, the lowest delivery flow to prevent clogging of the catheter.

### Drug delivery

KCC2 activity was blocked using VU0240551 (Sigma-Aldrich), VU0240551 was resuspended in DMSO as a 50mM stock solution, and later diluted in saline to the appropriate concentration. Recombinant human TrkB Fc chimera protein (TrkB-IgG; R&D company), used to chelate the endogenously released BDNF, was diluted in PBS. Parameters for drug delivery (dose, volume, time to reach maximal effect at the time of training) are consistent with previous reports (Gomez-Pinilla *et al.*, 2007; Austin and Delpire, 2011; Tashiro *et al.*, 2015). The specific delivery protocol and dosage are described in Fig. 1.

### Rehabilitation program

Beginning on day 5, exercised groups received a 60 minutes bicycling session. Animals were seated in a support harness with the hindlimbs hanging while their feet were secured to pedals with surgical tape. The custom-built motor-driven apparatus moves the hindlimbs through a complete range of motion during pedal rotation at a rate of 45 rpm (Houle *et al.*, 1999; Côté *et al.*, 2014). The rehabilitation program took place 5 days a week until completion of the study.

### Electrophysiological recordings and analysis

H-reflex recordings were performed 4 weeks post-SCI as described previously (Côté *et al.*, 2011; (2014). Rats were anaesthetized with ketamine (60 mg/kg) and xylazine (10 mg/kg) administered intraperitoneally. The tibial nerve was dissected free and mounted on a bipolar hook electrode for stimulation. Skin flaps were used to form a pool of mineral oil to prevent desiccation of the nerve. Bipolar wire electrodes (Cooner Wire) were inserted into the interosseus muscles for EMG recordings and the ground electrode inserted into the skin of the back. H-reflexes were evoked via an isolated pulse stimulator (A-M Systems) that delivered single bipolar pulses (100 μsec) to the tibial nerve, and H-and M-waves were recorded in the interosseus muscle in response to a range of increasing stimulus intensities. The intensity that elicited the maximal H-reflex amplitude (below the activation threshold for group Ib-II afferents ~1.2-1.4 MT) was then used to determine the properties of the M-wave and H-reflex as well as to evoke frequency-dependent depression (FDD). FDD was estimated using three series of 20 consecutive stimulations delivered at 0.3, 5, and 10 Hz. The experiment was first completed on the left leg, and the protocol then performed on the right leg.

EMG recordings were amplified (A-M system) and bandpass filtered (10–5 kHz), and the signal digitized (10 kHz) using a 1401 interface (Cambridge Electronic Design, CED) and fed to a computer running Signal 5 software (CED). Properties of the M-wave and H-reflex (n=8-13 animals/group, Table 1) are presented as mean ± SEM. For the analysis of the FDD, the first five responses to a train of stimulation were discarded to allow reflex stabilization. The 15 remaining responses were then averaged. Peak-to-peak amplitude of the M and H responses were measured and the H-reflex amplitude normalized to M_max_. The change in H-reflex peak-to-peak response at 5 Hz and 10 Hz was calculated as a percentage of the response obtained at 0.3 Hz. FDD data (n=8-13 animals/group) is presented as mean ± SEM.

After completion of the terminal experiment, rats were overdosed with Euthasol (390 mg/kg sodium pentobarbital and 50mg/kg phenytoin, i.p.) and the animal was either transcardiacally perfused with cold saline followed by 4% paraformaldehyde in PBS or fresh tissue was extracted for western blotting.

**Table 1.**
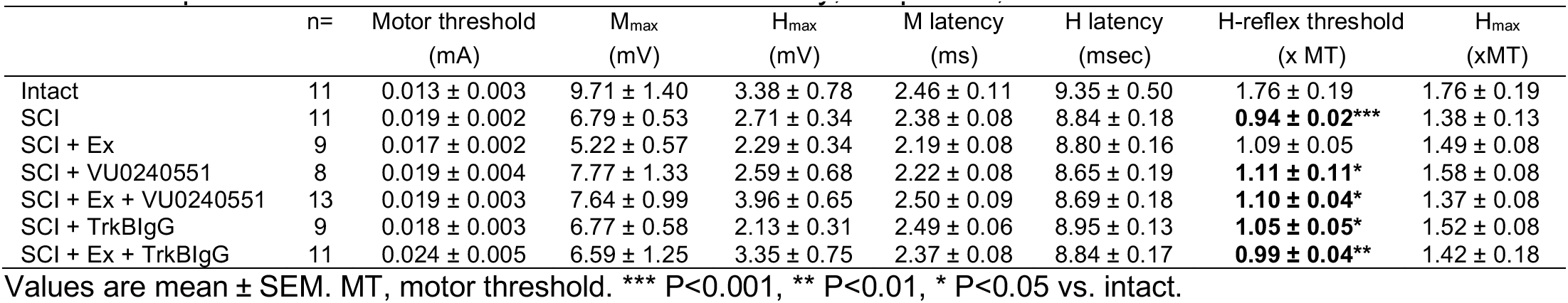
Properties of the M-Wave and H-Reflex: Latency, Amplitude, and Threshold

### Extraction of fresh tissue and Western blotting

Blocks of spinal tissue (L4-L5) were lysed in modified RIPA buffer (50mM Tris buffer pH 6.8, 1% Triton-X, 0.1% SDS, 1mM DTT, 0.5% deoxycholate, 150mM NaCl) containing protease and phosphatase inhibitors (Roche), 2mM PMSF and 1mM NaF. Protein concentration was determined using a BCA protein assay kit (Pierce). Samples (30μm of total protein) were subjected to SDS-PAGE (10-16% gels for BDNF, 4-20% gradient TGX gels (Biorad) for KCC2/NKCC1). Antibodies were targeted against KCC2 (1:1000, Millipore), BDNF (1:1000; Santa Cruz) or mouse T4 monoclonal antibody against NKCC (1:1000, Developmental Studies Hybridoma Bank), followed by incubation with HRP-conjugated secondary antibody (Jackson ImmunoResearch). Optical densities of protein bands were performed using Image J. Values for each sample were averaged, normalized to actin and combined for each group. Western blot data (n=6-11/group) is presented as a ratio of the intact group.

### Tissue preparation and immunohistochemistry

Spinal cord tissue was post-fixed overnight at 4°C, transferred to 30% sucrose for cryoprotection and cut transversally (40 μm sections) using a cryostat. Free floating sections were permeated in PBS containing 5% donkey serum and 0.1% Triton X-100 for one hour at room temperature and incubated with antibodies targeted against KCC2 (1:1000; Millipore) and ChAT (1:100, Millipore) for 24h. After brief washes in PBS, spinal cord sections were incubated in secondary antibody conjugated to FITC and rhodamine (1:400; Jackson Immunoresearch) for two hours at room temperature. Stacked images of motoneurons (40μm sections) were acquired with a spectral confocal & multiphoton system (Leica TCS SP2). The fluorescence intensity of KCC2 immunolabelling on the plasma membrane of motoneurons (identified by ChAT^+^, typical large size, and location within the ventral horn) was measured by averaging the integrated area of the density curve obtained by drawing three lines across each motoneurons (six data points) using ImageJ software (Côté *et al.*, 2014). A minimum of three lumbar motoneurons were averaged per animal (n=5-8 animals/group).

### Statistical analysis

Significant differences for the properties of the M-wave and H-reflex as well as immunoblotting data were determined by either using a one-way ANOVA followed by the Holm-Sidak or a Kruskal-Wallis one-way ANOVA on ranks followed by Dunn’s method if the sample variables did not fit a normal distribution (Shapiro-Wilk) or were not equally variant (Brown-Forsythes). Significant effects on the amplitude of the H-reflex by stimulation frequency and treatment group for the FDD were determined by a two-way ANOVA followed by the Holm-Sidak *post-hoc* test and the possible interaction of these factors with the variable was determined. Linear regression analysis was used to correlate NKCC1 to KCC2 and KCC2 to BDNF protein levels obtained by western blotting. All data are presented as mean ± SEM. Statistical analysis was performed using Sigma Plot software 14.0 and statistically significant levels were set to P<0.05.

## Results

### KCC2 activity is required for H-reflex recovery after chronic SCI

SCI induces a reduction in KCC2 expression in the lumbar enlargement of the spinal cord that is associated with the establishment and maintenance of spasticity after SCI (Boulenguez *et al.*, 2010). Our earlier findings illustrate that motor training prevents the injury-induced decrease in KCC2 expression on lumbar motoneurons and reduces spasticity (Côté *et al.*, 2014), but a causal relationship remained to be established. We therefore hypothesized that the decrease in hyperreflexia triggered by exercise requires KCC2 activity. We developed an intrathecal catheter delivery method to assess the effect of pharmacological inhibition of KCC2 on activity-dependent H-reflex recovery after SCI. KCC2 activation was specifically and only prevented during the daily rehabilitation session for 4 weeks post-SCI (Fig. 1B) at which time the properties of the H-reflex as well as its modulation was assessed. Because of its monosynaptic nature, the location of the neuroplasticity triggered by the injury and or inhibition is limited to primary afferent from group Ia fibers, α-motoneuron and their synaptic connection. In response to a tibial nerve stimulation, two successive responses can be recorded from the interosseous muscle, i.e. the M-wave results from the direct activation of motor axons while the H-reflex is evoked by the activation of Ia afferents that form a monosynaptic connection with motoneurons (Fig. 2A).

**Figure 2.**
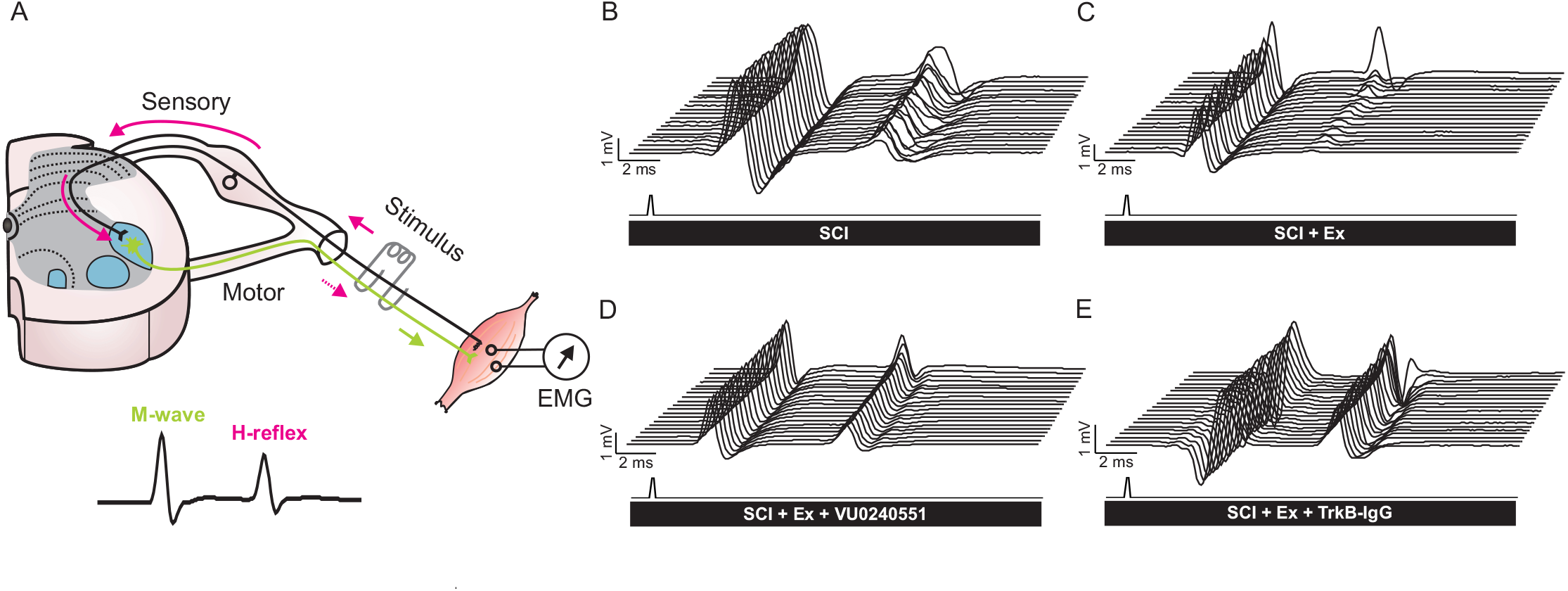
Representative recordings of H-reflexes evoked by a train of stimulation to the tibial nerve in the interosseus muscle. A) The stimulation of the tibial nerve evokes a volley in Ia afferents (solid pink arrows) that monosynaptically excite alpha motoneurons. The M-wave (green arrow) precedes the H-reflex (dotted pink arrow) and is due to the direct activation of motor axons. B-E) Typical EMG recordings over a series of 20 stimulations to the tibial nerve illustrating that during a 10Hz stimulation train, the depression of the H-reflex is impaired after SCI (B) but substantially restored in exercised animals (C). However, the exercised groups that received VU0240551 (D) or TrkB-IgG (E) during the daily rehabilitation session exhibited a very modest depression as compared to exercised animals. Overall, blocking KCC2 or BDNF activity in exercised animals (D-E) yields responses similar to non-exercised SCI (B).

The recruitment curve was used to determine the amplitude of the maximal m-wave (M_max_), the response of all motor units with supramaximal stimulation of axons of the tibial nerve, and of the maximal H-reflex amplitude (H_max_). To assess the relative proportion of motoneurons recruited through the monosynaptic reflex loop *vs.* the activation of the entire motor pool, the H_max_/M_max_ ratio was also calculated. The H_max_/M ratio was also used to evaluate the relative activation of the motor pool required to reach maximal reflex amplitude. Neither SCI nor VU0240551 affected the M-wave or H-reflex latency, the amplitude of the maximal M-wave (M_max_) and H-reflex (H_max_), the stimulation intensity at which H_max_ is evoked and the stimulation threshold to evoke a M-wave (Table 1). However, rehabilitation prevented the decrease in the stimulation intensity required to evoke a H-reflex after SCI. More importantly, blocking KCC2 activity prevented the effect of exercise with H-reflex thresholds similar to unexercised SCI but significantly lower than intact. Together, these results suggest that the decrease in H-reflex excitability triggered by exercise after a chronic SCI requires KCC2 activity.

After SCI, the excitability of the spinal cord gradually increases as a result of the injury. The transition to a state of hyperreflexia is well established by one-month after SCI (Bennett *et al.*, 1999; Yates *et al.*, 2008) and the reduction in the low frequency-dependent depression (FDD) of the H-reflex widely accepted as a reliable correlate of spasticity (Thompson *et al.*, 1992; Grey *et al.*, 2008; Boulenguez *et al.*, 2010). The improvement in FDD observed in exercised animals after SCI is characterized by the presence of a clear depression following a series if stimulations at 10Hz (Fig. 2C) while the depth of the modulation is much shallower in an animal not undergoing a rehabilitation program (Fig.2B). VU0240551 prevented the activity-dependent recovery triggered by exercise and the FDD was very modest FDD (Fig.2D). If some level of H-reflex modulation is still present after SCI regardless of exercise or VU0240551, as all groups displayed a decrease in H-reflex amplitude as the stimulation frequency increased, there was a remarkable difference in the depth of modulation across groups (Fig.3A-D). Figure 3E shows the averaged FDD for each group at each frequency expressed as a percentage of the response at 0.3Hz, a frequency at which there is no or little depression of the H-reflex. Higher H-reflex amplitude depicts less depression of the reflex. A two-way ANOVA revealed statistically significant differences across stimulation frequency (P<0.001) and across experimental groups (P<0.001) with a significant interaction between frequency and groups (P<0.001). *Post-hoc* comparisons suggested that 5 and 10 Hz values were different from 0.3Hz in all groups (P<0.001). The FDD drastically decreased after SCI with the amplitude of the H-reflex increasing from 13±3% to 57±7% at 5Hz and from 6±2% to 46±6% at 10Hz. Rehabilitation prevented this decrease with values not significantly different from uninjured animals at 5Hz and 10Hz (24±2% and 6±2% respectively), but considerably lower (i.e. more depression) than SCI. Blocking KCC2 prevented the recovery of FDD observed in exercised animals. VU0240551 significantly decreased the depression observed in exercised animals and had a FDD similar to SCI at 5Hz and 10Hz (47±6% and 40±6%). These results suggest that exercise failed to return H-reflex modulation after a chronic SCI when KCC2 cannot be activated during training.

**Figure 3.**
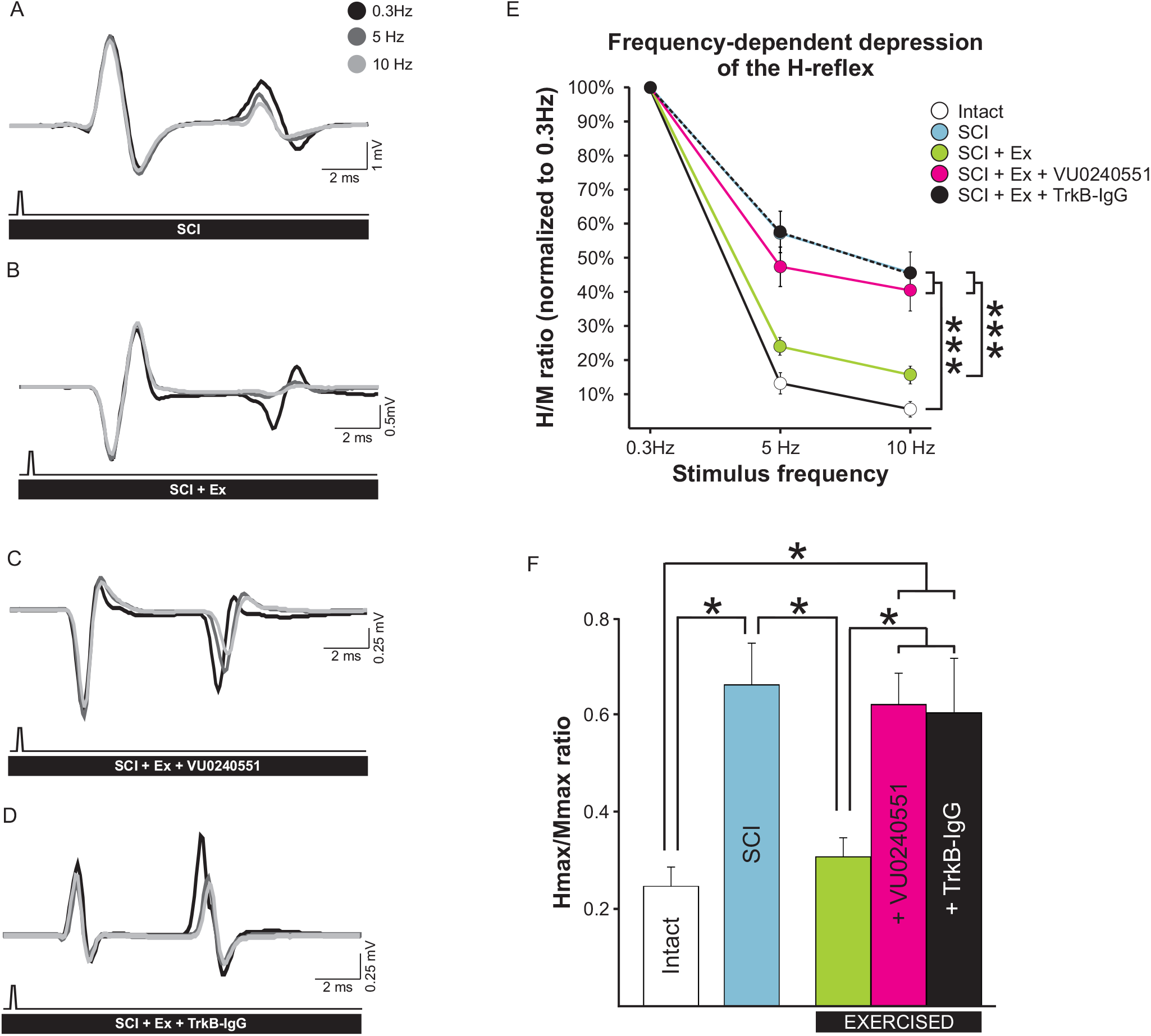
The activity-dependent recovery of the FDD in exercised animals is prevented by blocking KCC2 activity or scavenging BDNF after SCI. ***A-D)*** Averaged H-reflex traces (n>15) evoked by the stimulation of the tibial nerve at 0.3Hz (black), 5Hz (dark grey), 10Hz (light grey). Increasing stimulus frequency from 0.3Hz to 10Hz decreased H-reflex amplitude in all animal groups. ***E)*** There was a statistically significant difference across stimulation frequency (p<0.001) and across experimental groups (p<0.001) and the interaction between frequency and groups was also significant (p<0.001). H-reflexes were significantly smaller at 5Hz and 10 Hz as compared to 0.3 Hz in all groups. The FDD drastically decreased after SCI (blue, see also A) with the amplitude of the H-reflex increasing from 13±3% in intact animals to 57±7% in SCI at 5Hz (P<0.001) and from 6±2% to 46±6% at 10Hz (P<0.001). Rehabilitation prevented this decrease with values not significantly different from uninjured animals at 5Hz (24±2%; P=0.358) and 10Hz (6±2%; P=0.653)(green, see also B), but displayed considerably more depression than SCI at both stimulation frequencies (P<0.001). VU0240551 prevented the recovery of FDD (pink, see also C) observed in exercised animals. SCI+Ex+VU0240551 had FDDs similar to SCI at 5Hz (47±6%; P=0.317) and 10Hz (40±6%; P=0.650), but the depression was significantly lower than in exercised animals in which KCC2 activity was not blocked (P=0.002 and P<0.001 respectively). Scavenging BDNF also prevented the activity-dependent recovery of the FDD (black, see also D) at 5Hz and 10Hz (P<0.001) with values similar to sedentary SCI controls at 5Hz (P=0.951) and 10Hz (P=0.986) and SCI+Ex+VU0240551 (P=0.309 and P=0.778, respectively). Data is presented as mean ± SEM expressed as a percentage of the response obtained at 0.3Hz. *** P<0.001. Twoway ANOVA analysis with Holm-Sidak *post hoc* test, n=9-11 animals/group. ***F)*** The H_max_/M_max_ ratio, was also significantly different across groups (P<0.001). *Post-hoc* analysis revealed that SCI increases the H_max_/M_max_ ratio (P<0.027). Exercise restored H_max_/M_max_ which was lower than SCI values (P=0.045) but similar to intact (P=0.986). VU0240551 prevented the activity-dependent decrease in the H_max_/M_max_ which were not different from SCI (P=0.980) but significantly higher than exercised SCI (P=0.041) and intact (P=0.021). Scavenging BDNF also prevented the activity-dependent decrease in the H_max_/M_max_ ratio (P=0.033) with values similar to SCI (P=0.974) and SCI+Ex+VU0240551 (P=0.858). For illustration purposes, significance is illustrated for 10Hz only. H_max_/M_max_ ratios are presented as mean ± SEM. *P<0.05. One-way ANOVA analysis with Holm-Sidak *post hoc* test, n=9-11 animals/group).

The H_max_/M_max_ ratio, which estimates the fraction of motoneurons recruited relative to the activation of the entire motor pool, and is recognized as an estimate of motoneuronal excitability, was also significantly different across groups (P<0.001). Blocking KCC2 during rehabilitation sessions prevented the activity-dependent decrease in the H_max_/M_max_ ratio with SCI+Ex+VU0240551 animals displaying similar values to SCI but significantly higher than exercised and intact (Fig. 3F). This further support that KCC2 activity is required for the activity-dependent decrease in spinal hyperexcitability.

### BDNF activity is required for H-reflex recovery after chronic SCI

Because BDNF-TrkB signaling is known as a potent regulator of KCC2 in neurons (Rivera *et al.*, 2002; 2004; Coull *et al.*, 2005; Ferrini and De Koninck, 2013), we sought to further consider a causal relationship between the upregulation of BDNF and KCC2. If such a link exists after chronic SCI, preventing BDNF action, specifically during the training session, should prevent both the increase in KCC2 expression and the exercise-dependent recovery of the H-reflex modulation. We first examined the effect of the BDNF-sequestering TrkB-IgG on H-reflex properties.

Similar to what we observed when blocking KCC2 activity, chelating endogenous BDNF did not modify any features of the H-reflex but for the stimulation threshold to initiate a response (Table 1). TrkB-IgG prevented rehabilitation to restore the stimulation intensity required to initiate a H-reflex close to intact levels and this group displayed values similar to the SCI group and significantly lower than intact. This suggests that the decrease in H-reflex excitability triggered by exercise after SCI not only requires KCC2 activity, but also BDNF signaling. Scavenging BDNF also prevented the activity-dependent decrease in the H_max_/M_max_ ratio with values similar to SCI and SCI+Ex+VU0240551 (Fig.3F).

Similar to VU0240551 (Fig. 2D), TrkB-IgG prevented the activity-dependent recovery triggered by exercise and the FDD was very modest FDD (Fig. 2E and 3D). Overall, scavenging BDNF significantly prevented the activity-dependent recovery of the FDD (Fig. 3E) (5 and 10Hz, P<0.001) and H-reflex amplitude values were no different from unexercised SCI or exercised animals that had received VU0240551. This provides evidence that rehabilitation fails to reduce hyperexcitability in spinal networks after SCI when BDNF activity was prevented. Blocking BDNF or KCC2 activity yielded very comparable results.

### Blocking KCC2 or scavenging BDNF after SCI does not affect hyperreflexia in unexercised animals

To exclude any confounding factor and verify that the results observed are not simply due to the effect of the drugs on spinal networks rather than blocking the exercise-dependent recovery of the H-reflex modulation, we assessed if there was any effect of blocking KCC2 or BDNF activity in unexercised SCI rats. When comparing SCI animals that received VU0240551 or TrkB-IgG to animals that received the vehicle, none of the general features of the M-wave and H-reflex were significantly altered 4 weeks post-SCI (Table 1). These results suggest that, at this dose, drugs alone had no effect on spinal excitability when no rehabilitation program was implemented. Similarly, the modulation of the H-reflex was not different after SCI whether a drug treatment was used or not (*not shown*). The FDD of SCI animals that received VU0240551 or TrkB-IgG was not statistically different from SCI animals receiving the vehicle, with all groups displaying a very modest depression at both 5Hz (respectively 41±5%, 47±6, 57±7% %) and 10Hz and (30±5%, 32±5%, 46±6%). This confirms that blocking KCC2 or BDNF activity had no effect after SCI in untrained animals when KCC2 and BDNF levels are already low in the spinal cord and suggests that the drug itself does not impact motor recovery. No further analysis was carried out on these control groups as the objective of the study is to investigate if KCC2 is critical to the beneficial effect of activity-based therapies.

### BDNF signaling is required for the activity-dependent increase in KCC2 expression

We then performed western blot analysis on tissue from the lumbar enlargement to evaluate the effect of blocking KCC2 or BDNF activity. One-way ANOVA confirmed differences in KCC2 protein expression across groups (P<0.001). Further *post hoc* analysis showed that KCC2 expression is modulated after SCI, decreasing to 64±3% of intact values (Fig. 4A, see also Côté *et al.*, 2014). Interestingly, the injury-induced downregulation in KCC2 expression was prevented by the rehabilitation program, whether KCC2 activity was blocked or not, with KCC2 expression levels not different from intact. This suggests that VU0240551 only affected KCC2 activity, but did not affect protein synthesis. On the contrary, scavenging BDNF prevented the activity-dependent increase in KCC2 expression with values similar to unexercised animals (63 ± 5%) and lower than intact or exercised animals with, suggesting that BDNF activity is required to restore KCC2 expression induced by exercise.

One-way ANOVA also confirmed a difference in BDNF protein expression between groups (P<0.001). Exercise prevents the decrease in BDNF expression after a chronic SCI (Fig. 4B). Animals that followed a rehabilitation program had higher BDNF expression levels as compared to unexercised SCI, and scavenging BDNF during training yielded lower levels. Surprisingly, blocking KCC2 activity prevented the beneficial effect of exercise on BDNF levels with values significantly lower than exercised animals receiving the vehicle, suggesting a mechanism by which KCC2 can regulate BDNF levels.

**Figure 4.**
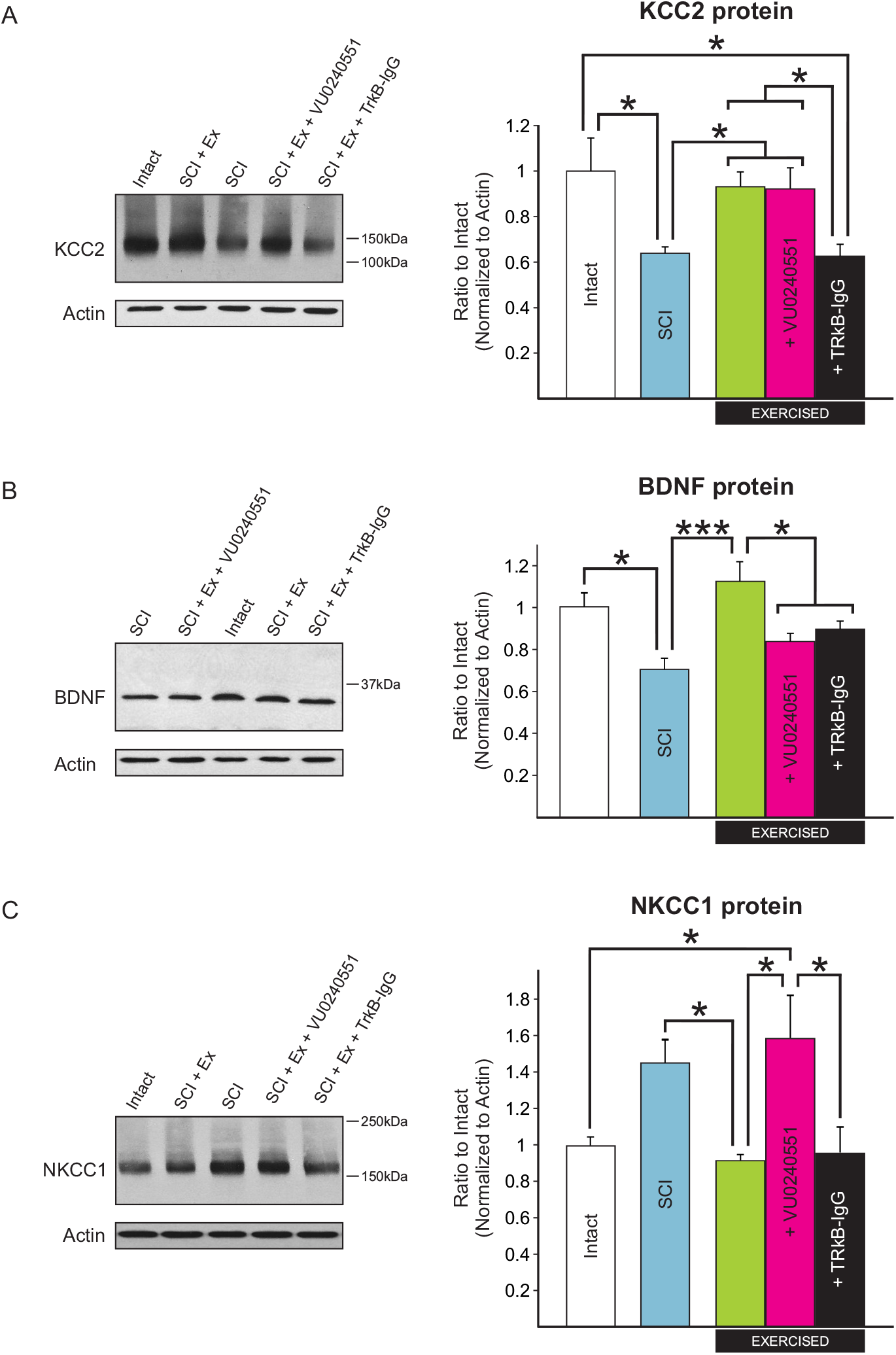
The activity-dependent modulation of KCC2, NKCC1 and BDNF protein expression in the lumbar enlargement of SCI rats requires KCC2 and BDNF activity. Western blot analysis shows that **A)** One-way ANOVA confirmed differences in KCC2 protein expression across groups (P<0.001). Further *post hoc* analysis showed that KCC2 expression is modulated after SCI, decreasing to 64±3% of intact values (P=0.015). This decrease was prevented by a rehabilitation program whether KCC2 was blocked (P<0.034) or not (P<0.021) as both SCI-Ex and SCI+Ex+VU0240551 displayed KCC2 expression levels similar to intact (93 ± 7%, P=0.898 and 92 ± 10%, P=0.944 respectively). Scavenging BDNF prevented the activity-dependent increase in KCC2 expression with values similar to sedentary animals (63 ± 5%, P=0.989), and lower than intact (P=0.015) and SCI+Ex (P=0.022). One-way ANOVA analysis (P<0.001) with Holm-Sidak *post hoc* test, n = 6-11/group). **B)** One-way ANOVA confirmed a difference in BDNF protein expression between groups (P<0.001). The decrease in BDNF expression after a chronic SCI as compared to intact (P=0.024) was prevented by exercise. SCI-Ex displayed BDNF expression levels higher than SCI (P<0.001) and similar to intact (P=0.425). Blocking KCC2 activity prevented the beneficial effect of exercise on BDNF levels with values significantly lower than exercised animals receiving the vehicle (P=0.011). Scavenging BDNF during training yielded lower levels of BDNF in the lumbar enlargement (P=0.043). **C)** NKCC1 protein expression in the lumbar enlargement was significantly different between groups (P=0.003). NKCC1 levels tended to increase after SCI. Although this increase did not reach statistical significance (P=0.068), rehabilitation returned NKCC1 expression levels close to intact (93 ± 4%, P=0.964) with significantly less NKCC1 in the lumbar enlargement than sedentary SCI animals (147 ± 14%, P=0.036). Blocking KCC2 prevented the activity-dependent decrease in NKCC1 expression and SCI+Ex+VU0240551 displayed values similar to sedentary SCI animals (161 ± 24%, P=0.930) but higher than SCI-Ex (P=0.012). Scavenging BDNF did not prevent the activity-dependent decrease in NKCC1 expression (97 ± 16%, P=0.969). One-way ANOVA analysis with Holm-Sidak *post hoc* test, n = 6-11/group). One-way ANOVA analysis (P<0.001) with Holm-Sidak *post hoc* test, n = 6-11/group). Protein levels are presented as mean ± SEM. *P<0.05, ***P<0.001 vs intact.

A regression analysis showed that there is a significant linear relationship between BDNF and KCC2 expression in the lumbar enlargement of the spinal cord (Fig.5A). Data from SCI that received the vehicle or TrkB-IgG are clustered in the lower left quadrant with low expression levels of both BDNF and KCC2 while intact and SCI animals that followed a rehabilitation program are grouped in the top right quadrant displaying higher levels of both proteins. This further supports a role for BDNF in regulating KCC2.

**Figure 5.**
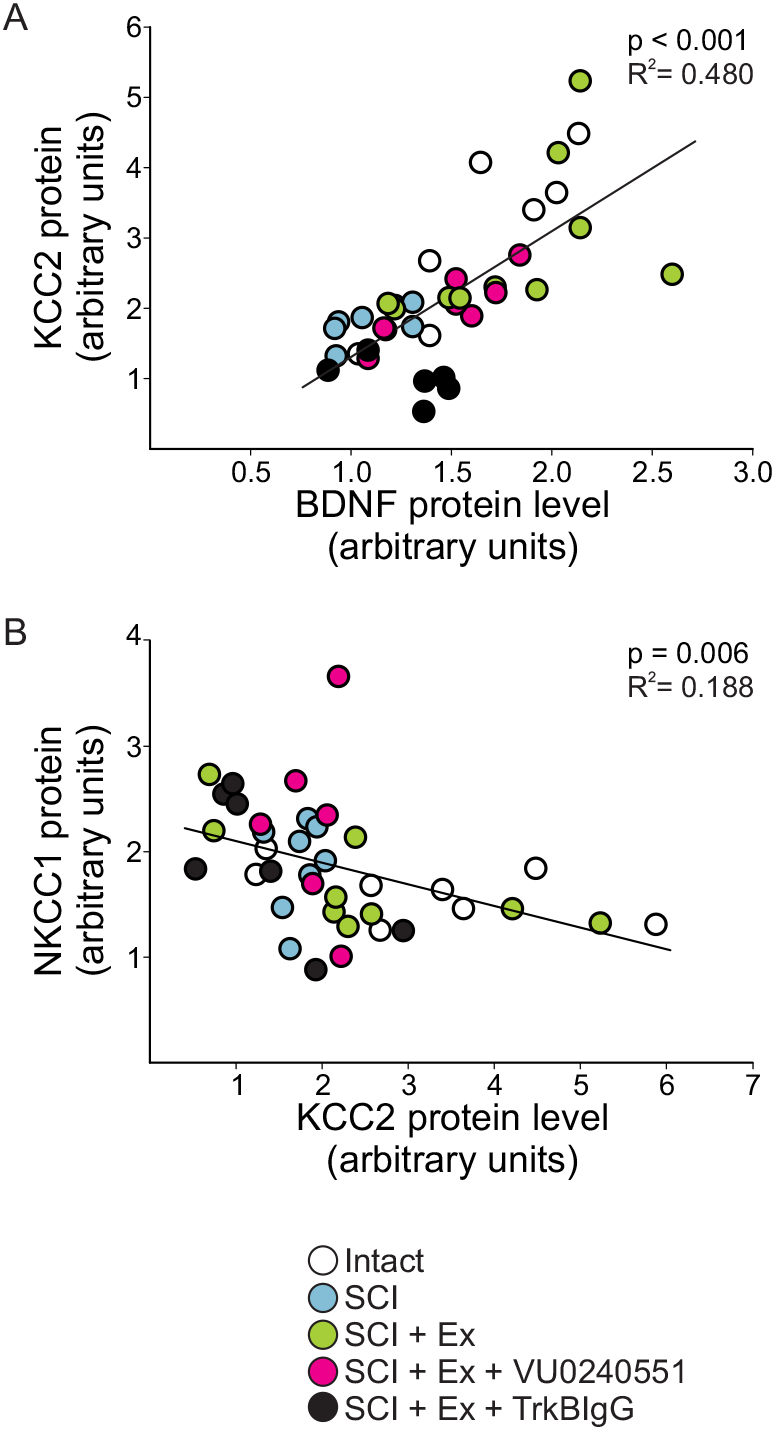
Relationship between the expression of NKCC1, KCC2 and BDNF in the lumbar spinal cord. ***A)*** A linear regression analysis showed that there is a significant positive relationship between BDNF and KCC2 expression level in the lumbar spinal cord (P<0.001, R^2^=0.480). SCI and SCI+Ex+TrkB-IgG groups are clustered in the lower left quadrant with low expression levels of BDNF and KCC2 while intact and SCI+Ex are grouped in the top right quadrant displaying higher levels of both proteins. ***B)*** A linear regression analysis illustrate a significant negative relationship between NKCC1 and KCC2 expression level in the lumbar spinal cord (P=0.006, R^2^=0.188) with low levels of KCC2 expression associated to higher levels of NKCC1.

Although the shift in chloride homeostasis and impaired FDD after chronic SCI was shown to depend on a decrease in KCC2 expression (Boulenguez *et al.*, 2010), the inwardly directed Na^+^-K^+^-Cl^-^ cotransporter isoform 1, NKCC1, is also upregulated after SCI (Cramer *et al.*, 2008; Hasbargen *et al.*, 2010; Lee *et al.*, 2014). The relative expression of KCC2 and NKCC1 critically determine [Cl^-^]_i_and GABA-mediated responses. NKCC1 is responsible to bring Cl^-^ in the cell, in opposition to KCC2, so that a simultaneous decrease in KCC2 and increase in NKCC1 after SCI (Coté *et al.*, 2014) would act synergistically to depolarize E_cl-_. We therefore measured changes in NKCC1 protein expression in the lumbar enlargement and found a significant difference between groups (Fig. 4C, P=0.003). Rehabilitation returned NKCC1 expression levels close to intact values (93 ± 4%) with significantly less NKCC1 in the lumbar enlargement than unexercised SCI animals (147 ± 14%). VU0240551 prevented the restoration of NKCC1 expression levels following a rehabilitation program with values not different from unexercised SCI animals (161 ± 24%) but higher than exercised. Scavenging BDNF did not affect NKCC1 expression (97 ± 16%). This suggests that NKCC1 is also modulated in an activity-dependent manner after SCI and may contribute to the shift in chloride homeostasis induced by exercise. A regression analysis was also carried out to address the expression of NKCC1 in the lumbar spinal cord as a function of KCC2. We observed a significant negative relationship between NKCC1 and KCC2 expression levels in the lumbar spinal cord (Fig.5B). Consistent with their opposing directionality in chloride transport, this supports previous findings suggesting that NKCC1 and KCC2 protein levels are reciprocally regulated.

As hyperreflexia and spasticity are specifically associated with a KCC2 increase in motoneuronal membrane in the lumbar enlargement (Boulenguez *et al.*, 2010), we sought to investigate if the BDNF dependent increase in KCC2 expression involved in activity-dependent recovery occurs at this specific location. In the ventrolateral spinal cord, KCC2 labelling is particularly strong around the motoneuronal membrane (Fig. 6). Exercise increased KCC2 expression around the motoneuronal membrane and less cytoplasmic clusters are visible in the soma, suggesting a decrease in the internalization of KCC2 into vesicles. Blocking KCC2 or BDNF activity during exercise prevented this effect, i.e. the activity-dependent restoration of KCC2 around the motoneuronal membrane was not observed.

**Figure 6.**
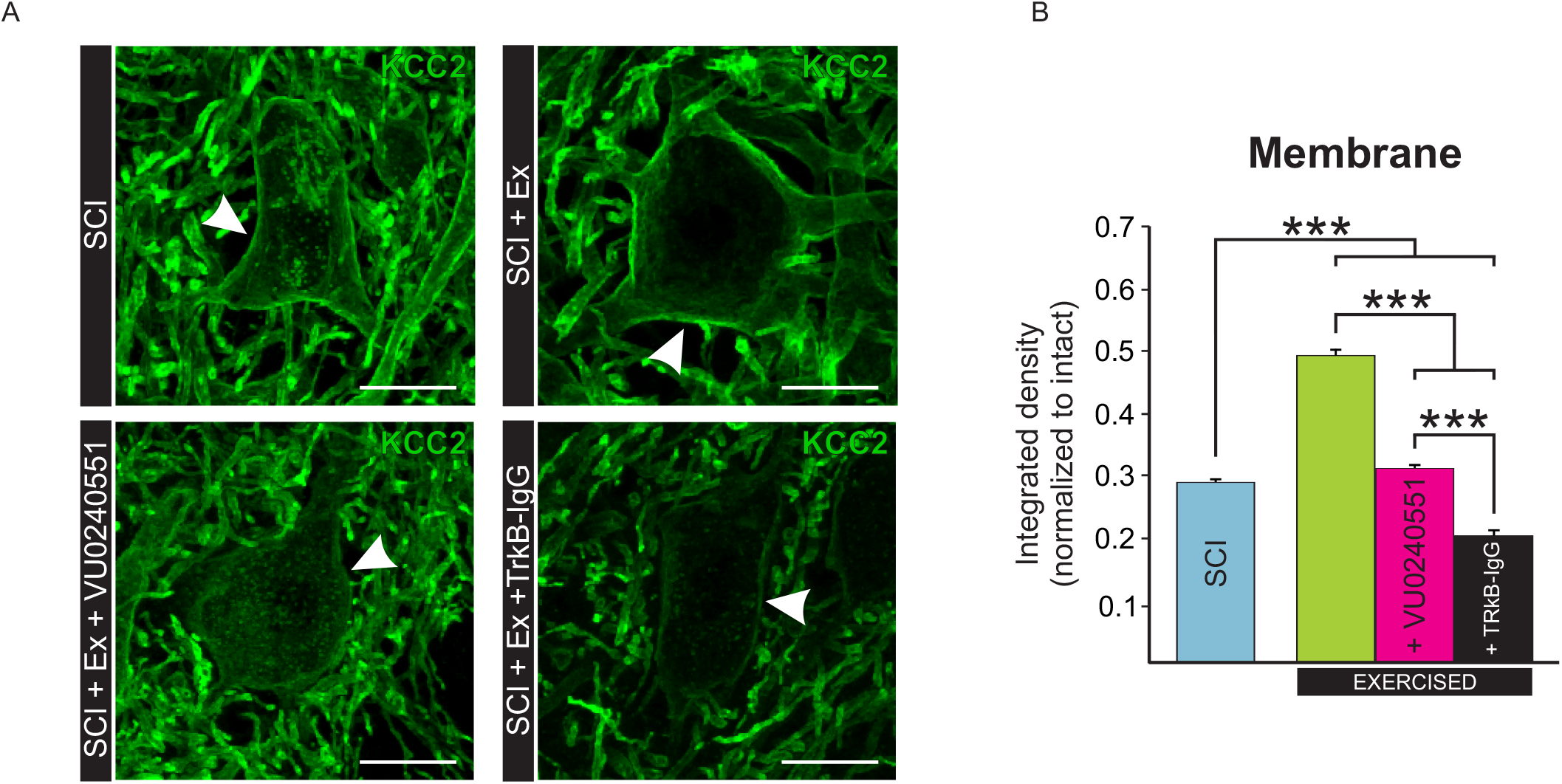
Blocking KCC2 or BDNF activity during exercise prevents the activity-dependent increase in KCC2 levels in lumbar motoneuronal membrane. ***A)*** Digital images showing KCC2 expression in the membrane around the motoneuronal soma (white arrows) and also in dendrites in the ventral horn of the lumbar enlargement. ***B)*** Quantification of the integrated density of KCC2 around the motoneuronal membrane reveals that exercise increases KCC2 immunoreactivity around the membrane (P<0.001) and that blocking KCC2 or BDNF activity prevents this activity-dependent increase after SCI (P<0.001). Data are presented as means ± SEM. Kruskal-Wallis ANOVA on ranks (p<0.001) followed by Dunn’s method, n=5-8 animals per group. ***P<0.001, Scale bar = 25 μm.

## Discussion

Rehabilitation approaches that promote repetitive motor activity are widely used in the clinic and are a critical component of successful functional recovery in SCI individuals. A variety of exercise programs have revealed potential to alleviate spasticity, but the molecular pathways involved in functional recovery remains elusive. Here, we show for the first time that 1) KCC2 activity is required to the beneficial effect of exercise on spasticity; 2) this activity-dependent increase in KCC2 levels is regulated by BDNF.

### KCC2 activity is required to decrease spasticity after chronic SCI

Under physiological conditions, KCC2 extrudes Cl^-^ from neurons maintaining low levels of [Cl^-^]. This insures that GABAergic synaptic transmission remains hyperpolarizing as Cl^-^ flows inwardly along its electrochemical gradient. A condition that negatively affects KCC2 function leads to an accumulation of [Cl^-^]_i_which undermines GABA_A_-mediated inhibition (Rivera *et al.*, 1999; Hübner *et al.*, 2001; Coull *et al.*, 2003). In spinal motoneurons, genetic or pharmacological manipulations that decrease KCC2 expression/function elicit a ~10-20mV depolarizing shift in E_cl_-which increases neuronal excitability and impairs postsynaptic inhibition (Hübner *et al.*, 2001; Jean-Xavier *et al.*, 2006; Bos *et al.*, 2013). Similarly, the SCI-induced decrease in motoneuronal KCC2 expression and subsequent shift in chloride homeostasis also contributes to spinal hyperexcitability and spasticity (Boulenguez *et al.*, 2010; Mòdol *et al.*, 2014). SCI, as other CNS pathologies, recapitulate developmental programs with low levels of KCC2 and consequent depolarizing shift in E_GABAA_ reminiscent of immature neurons (reviewed in Kaila *et al.*, 2014a).

After chronic SCI, the loss of supraspinal control and increase in the gain of afferent feedback increases the excitability of spinal reflexes. This results in permanent undesirable effects such as inappropriate timing of activation of muscles during movement and failure to adjust reflex excitability to meet task requirements. Rehabilitation programs can both restore KCC2 expression on lumbar motoneurons and improve reflex recovery (Côté *et al.*, 2014), but the causal relationship linking the shift in chloride homeostasis and the recovery of motor function remains unclear. Here, we have used a chronic delivery of VU0240551, a highly specific KCC2 blocker (Delpire *et al.*, 2009), to evaluate the functional role of KCC2 in motor recovery after SCI. We insured that its effect was specifically targeted to KCC2 function during rehabilitation (vs. spontaneous recovery) with a delivery protocol restricted to the exercise period and by taking advantage of its relatively short therapeutic window (Austin and Delpire, 2011).

We have used the FDD, which is indicative of the function of the spinal inhibitory system, is dependent on GABA_A_ receptor-mediated inhibition in rodents and is associated to spasticity (Jolivalt *et al.*, 2008; Boulenguez *et al.*, 2010; Lee-Kubli and Calcutt, 2014). The lack of significant effect of VU0240551 in unexercised SCI animals suggests a low target availability and confirms that our observations are not solely relying on the effect of the drug, but to its interaction with activity-based therapy. While blocking KCC2 with VU0240551 did not affect KCC2 expression in the lumbar enlargement, confocal analysis distinctly illustrated a decrease expression on motoneuronal membrane. This supports findings suggesting that VU0240551 mainly affects post-translational mechanisms (vs. protein synthesis), including preventing shuttling to the membrane, which is also supported by the increased presence of clusters in the cytoplasm (Fig. 6A).

In agreement with our earlier studies, animals that underwent a rehabilitation program after SCI displayed much less spinal hyperexcitability (Côté *et al.*, 2011; 2014). VU0240551 prevented the activity-dependent recovery of reflex modulation, indicating that the beneficial effect of exercise is critically dependent on KCC2 activity. Together, our data suggests that rehabilitation increases both KCC2 synthesis (Fig. 4A) and post-translational mechanisms responsible for KCC2 trafficking to the membrane (Fig., 6A). Such mechanisms include increased insertion rate to the membrane and/or decreased internalization by endocytosis (Lee *et al.*, 2007).

### Activity-dependent modulation of KCC2 protein is BDNF-dependent

KCC2 is dynamically modulated by multiple intra-and intercellular signaling pathways (Fiumelli and Woodin, 2007), with the most prevalent being BDNF signaling onto TrkB receptors (TrkB; Rivera *et al.*, 2002; 2004; Coull *et al.*, 2005; Ferrini and De Koninck, 2013). The polarity of BDNF-dependent regulation of KCC2 expression differs depending on the developmental stage and the integrity of the CNS. While BDNF reduces GABAergic inhibition through KCC2 downregulation in mature neurons (Rivera *et al.*, 2002; 2004; Coull *et al.*, 2005; Boulenguez *et al.*, 2010; Huang *et al.*, 2017), it promotes an increase in KCC2 expression in embryonic neurons or following an axotomy in the adult (Aguado *et al.*, 2003; Payne *et al.*, 2003; Carmona *et al.*, 2006).

After a chronic SCI, activity-based therapies increase KCC2 and BDNF levels in the lumbar spinal cord (Côté *et al.*, 2014; Chopek *et al.*, 2015; Tashiro *et al.*, 2015). Increased BDNF levels is associated with the normalization of motoneuronal properties (Beaumont *et al.*, 2008) and locomotor recovery (Boyce *et al.*, 2007; 2012; Ying *et al.*, 2008), and is positively correlated to the restoration of reflex modulation, suggesting a linear relationship with hyperreflexia (Côté *et al.*, 2011). Interestingly, BDNF upregulates KCC2 in SDH neurons and attenuate central sensitization through reinstating GABAergic inhibition (Huie *et al.*, 2012; Huang *et al.*, 2017). In order to identify BDNF-TrkB as an activity-dependent regulator of KCC2 after SCI, we used a TrkB-IgG, a chimeric molecule comprised of the extracellular domain of TrkB that sequesters BDNF by competing with endogenous TrkB. TrkB-IgG delivered during training yielded lower levels of BDNF as compared to exercised animals that received the vehicle, confirming that TrkB-IgG treatment was effective at scavenging BDNF. Our results illustrate that depleting BDNF availability not only prevented the rehabilitation-induced recovery of the FDD, but more importantly the increase in KCC2 motoneuronal expression triggered by exercise. This indicate that BDNF contributes to decrease hyperreflexia after SCI through regulating KCC2 expression and restoring chloride homeostasis after SCI.

In the hippocampus, enhanced membrane insertion and retention of KCC2 depends on BDNF-TrkB signaling in response to increased network activity (Khirug *et al.*, 2010; Puskarjov *et al.*, 2015). The results presented here provide evidence that a similar mechanism is at play in the spinal cord. Chronic SCI recapitulates an earlier developmental state in which BDNF promotes KCC2 upregulation and restores GABA-mediated inhibition. Our results suggest that the activity-dependent activation of BDNF/TrkB alters chloride extrusion and the global inhibitory action of GABA_A_-mediated responses as depicted by the restoration of the FDD in exercised animals. Among several possibilities, TrkB activation can regulate calpains (Zadran *et al.*, 2010). Interestingly, calpain-dependent proteolytic cleavage of KCC2 alters its ability to extrude Cl^-^ in the hippocampus and spinal cord (Puskarjov *et al.*, 2012; Zhou *et al.*, 2012) and contributes to the development of spasticity after SCI (Brocard *et al.*, 2016; Plantier and Brocard, 2017). Whether activity-dependent plasticity affects this pathway in motoneurons remains to be determined.

It is also worth noting that not only does chelating BDNF decreases KCC2 expression (Fig. 4A), but blocking KCC2 also appears to decrease BDNF levels in exercised animals (Fig. 4B). This suggest the presence of a retroactive feedback loop through which KCC2 can also regulate BDNF expression.

### BDNF is not responsible for the exercise-induced reciprocal regulation of KCC2 and NKCC1

Activity-dependent plasticity triggered by exercise after SCI not only increases the expression of KCC2 but also decreases the expression of the chloride intruder NKCC1 (Fig. 4C, see also Côté *et al.*, 2014), as both transporters act in synergy to restore [Cl^-^]_i_. VU0240551 prevented the effect of exercise on both KCC2 and NKCC1, so that their expression levels in the lumbar spinal cord maintained a negative correlation. Because NKCC1 staining did not reveal strong enough staining on motoneuronal membrane, but is strongly expressed in small glial cells surrounding motoneurons (Kanaka *et al.*, 2001) as well as primary afferent terminal (Stil *et al.*, 2009), identifying with precision if its expression was specifically downregulated in motoneurons was not possible.

Scavenging BDNF did not affect NKCC1 expression, suggesting that BDNF is not responsible for the activity-dependent reciprocal regulation of KCC2 and NKCC1. NKCC1 and KCC2 activity is regulated in a reciprocal fashion through the WNK-SPAK/OSR1 pathway, i.e. WNKs inactivates KCC2 and activates NKCC1, providing a coordinated control over [Cl^-^]_i_(Kahle *et al.*, 2010, Alessi *et al.*, 2014). Decreasing WNK1 activity triggers a hyperpolarizing shift in GABA responses by enhancing KCC2-mediated Cl^−^ extrusion (Friedel *et al.*, 2015) and reduces allodynia and hyperalgesia in SDH neurons (Kahle *et al.*, 2016). The WNK1 pathway can be activated by lower levels of [Cl^-^]i, a hallmark of SCI. Anecdotical reports suggest the involvement of WNK-SPAK/OSR1 pathway in activating NKCC1 at the lesion site after SCI (Lee *et al.*, 2014) but warrants further studies. Whether activity-based therapies decrease WNK1 activity remains to be determined, but our results strongly support this possibility.

### Therapeutic significance

Available anti-spastic drugs such as baclofen, diazepam, tizanidine and botulinum toxin have serious side effects including sedation, dizziness and a deep, long-lasting depression of spinal excitability that significantly reduces muscle activity and interferes with motor recovery (Dario and Tomei, 2004; Adams and Hicks, 2005; Elbasiouny *et al.*, 2010; Angeli *et al.*, 2012). The hyperexcitability of spinal networks in SCI individuals often leads to spasticity and co-contraction of flexor and extensor muscles due to less inhibitory reflexes (Boorman *et al.*, 1996; Mazzocchio and Rossi, 1997; Katz, 1999; Harkema, 2008). Inhibitory synapses constitute about 30% of all synapses in the spinal cord and are critical for the optimal function of neural circuits by regulating oscillatory behavior of neuronal networks, contributing to multifaceted aspects of neuronal processing, and limiting the extent of excitatory activity when required. Given the role of KCC2 in regulating the strength of inhibitory synaptic transmission, facilitating neuronal Cl^-^ extrusion by directly targeting KCC2 has emerged as a promising alternative (Doyon *et al.*, 2013; Kahle *et al.*, 2014). This study illustrates for the first time that the beneficial effect of exercise on spinal hyperexcitability requires the activation of the BDNF-KCC2 pathway to maintain chloride homeostasis in motoneurons after SCI. While BDNF is effective in improving spasticity, longterm administration has shown serious therapeutic drawbacks (Weishaupt *et al.*, 2012; Fouad *et al.*, 2013).

Targeting Cl^-^ transport by activating KCC2 and enhancing chloride extrusion can modulate GABAergic transmission *in vivo* and decrease neuropathic pain and hyperalgesia (Gagnon *et al.*, 2013; Ferrini *et al.*, 2017). This critically identifies this pathway as a potential pharmacological therapeutic target to improve hyperreflexia after a chronic SCI for individuals with comorbidities that delays the onset of physical therapy. Acting directly on chloride homeostasis would help to restore endogenous inhibition rather than actively depress excitability.

## ACKNOWLEDGEMENTS

We are in debt to Emerita Professor Marion Murray and Dr. Michel Lemay for their significant criticisms, comments, input and exchange of ideas on earlier versions of this manuscript.

## References

Adams MM, Hicks AL. Spasticity after spinal cord injury. Spinal Cord 2005; 43: 577–86.

Aguado F, Carmona MA, Pozas E, Aguilo A, Martinez-Guijarro FJ, Alcantara S,et al. BDNF regulates spontaneous correlated activity at early developmental stages by increasing synaptogenesis and expression of the K^+^/Cl^-^ co-transporter KCC2. Development 2003; 130: 1267–80.

Alessi DR, Zhang J, Khanna A, Hochdorfer T, Shang Y, Kahle KT. The WNK-SPAK/OSR1 pathway: master regulator of cation-chloride cotransporters. Sci Signal 2014; 7: re3.

Angeli C, Ochsner J, Harkema S. Effects of chronic baclofen use on active movement in an individual with a spinal cord injury. Spinal Cord 2012; 50: 925–7.

Austin TM, Delpire E. Inhibition of KCC2 in mouse spinal cord neurons leads to hypersensitivity to thermal stimulation. Anesth Analg 2011; 113: 1509–15.

Beaumont E, Kaloustian S, Rousseau G, Cormery B. Training improves the electrophysiological properties of lumbar neurons and locomotion after thoracic spinal cord injury in rats. Neurosci Res 2008; 62: 147–54.

Bennett DJ, Gorassini M, Fouad K, Sanelli L, Han Y, Cheng J. Spasticity in rats with sacral spinal cord injury. J Neurotrauma 1999; 16: 69–84.

Boorman GI, Lee RG, Becker WJ, Windhorst UR. Impaired “natural reciprocal inhibition” in patients with spasticity due to incomplete spinal cord injury. Electroencephalogr Clin Neurophysiol 1996; 101: 84–92.

Bos R, Sadlaoud K, Boulenguez P, Buttigieg D, Liabeuf S, Brocard C,et al. Activation of 5-HT2A receptors upregulates the function of the neuronal K-Cl cotransporter KCC2. Proc Natl Acad Sci USA 2013; 110: 348–53.

Boulenguez P, Liabeuf S, Bos R, Bras H, Jean-Xavier C, Brocard C,et al. Down-regulation of the potassium-chloride cotransporter KCC2 contributes to spasticity after spinal cord injury. Nat Med 2010; 16: 302–7.

Boyce VS, Mendell LM. Neurotrophic factors in spinal cord injury. Handb Exp Pharmacol 2014a; 220: 443–60.

Boyce VS, Mendell LM. Neurotrophins and spinal circuit function. Front Neural Circuits 2014b; 8: 59.

Boyce VS, Park J, Gage FH, Mendell LM. Differential effects of brain-derived neurotrophic factor and neurotrophin-3 on hindlimb function in paraplegic rats. Eur J Neurosci 2012; 35: 221–32.

Boyce VS, Tumolo M, Fischer I, Murray M, Lemay MA. Neurotrophic factors promote and enhance locomotor recovery in untrained spinalized cats. J Neurophysiol 2007; 98: 1988–96.

Brocard C, Plantier V, Boulenguez P, Liabeuf S, Bouhadfane M, Viallat-Lieutaud A,et al. Cleavage of Na(+) channels by calpain increases persistent Na(+) current and promotes spasticity after spinal cord injury. Nat Med 2016; 22: 404–11.

Carmona MA, Pozas E, Martinez A, Espinosa-Parrilla JF, Soriano E, Aguado F. Age-dependent spontaneous hyperexcitability and impairment of GABAergic function in the hippocampus of mice lacking trkB. Cereb Cortex 2006; 16: 47–63.

Chopek JW, Sheppard PC, Gardiner K, Gardiner PF. Serotonin receptor and KCC2 gene expression in lumbar flexor and extensor motoneurons posttransection with and without passive cycling. J Neurophysiol 2015; 113: 1369–76.

Côté M-P, Azzam GA, Lemay MA, Zhukareva V, Houle JD. Activity-dependent increase in neurotrophic factors is associated with an enhanced modulation of spinal reflexes after spinal cord injury. J Neurotrauma 2011; 28: 299–309.

Côté M-P, Gandhi S, Zambrotta M, Houle JD. Exercise modulates chloride homeostasis after spinal cord injury. J Neurosci 2014; 34: 8976–87.

Coull JA, Beggs S, Boudreau D, Boivin D, Tsuda M, Inoue K,et al. BDNF from microglia causes the shift in neuronal anion gradient underlying neuropathic pain. Nature 2005; 438: 1017–21.

Coull JA, Boudreau D, Bachand K, Prescott SA, Nault F, Sik A,et al. Trans-synaptic shift in anion gradient in spinal lamina I neurons as a mechanism of neuropathic pain. Nature 2003; 424: 938–42.

Cramer SW, Baggott C, Cain J, Tilghman J, Allcock B, Miranpuri G,et al. The role of cation-dependent chloride transporters in neuropathic pain following spinal cord injury. Mol Pain 2008; 4: 36.

Dario A, Tomei G. A benefit-risk assessment of baclofen in severe spinal spasticity. Drug Saf 2004; 27: 799–818.

Delpire E. Cation-Chloride Cotransporters in Neuronal Communication. News Physiol Sci 2000; 15: 309–12.

Delpire E, Days E, Lewis LM, Mi D, Kim K, Lindsley CW,et al. Small-molecule screen identifies inhibitors of the neuronal K-Cl cotransporter KCC2. Proc Natl Acad Sci USA 2009; 106: 5383–8.

Dietz V (2001) Spinal cord lesion: effects of and prespectives for treatment. Neural Plast 8:8390.

Doyon N, Ferrini F, Gagnon M, De Koninc Y. Treating pathological pain: is KCC2 the key to the gate? Expert Rev Neurother 2013; 13: 469–71.

Ferrini F, De Koninck Y. Microglia control neuronal network excitability via BDNF signalling. Neural Plast 2013; 2013: 429815.

Ferrini F, Trang T, Mattioli TA, Laffray S, Del’Guidice T, Lorenzo LE, Castonguay A, Doyon N, Zhang W, Godin AG, Mohr D, Beggs S, Vandal K, Beaulieu JM, Cahill CM, Salter MW, De Koninck Y (2013) Morphine hyperalgesia gated through microglia-mediated disruption of neuronal Cl(-) homeostasis. Nat Neurosci 16:183–192.

Ferrini F, Lorenzo LE, Godin AG, Quang ML, De Koninck Y. Enhancing KCC2 function counteracts morphine-induced hyperalgesia. Sci Rep 2017; 7: 3870.

Fiumelli H, Woodin MA. Role of activity-dependent regulation of neuronal chloride homeostasis in development. Curr Opin Neurobiol 2007: 17: 81–6.

Fouad K, Bennett DJ, Vavrek R, Blesch A. Long-term viral brain-derived neurotrophic factor delivery promotes spasticity in rats with a cervical spinal cord hemisection. Front Neurol 2013; 4: 187.

Friedel P, Kahle KT, Zhang J, Hertz N, Pisella LI, Buhler E, Schaller F, Duan J, Khanna AR, Bishop PN, Shokat KM, Medina I (2015) WNK1-regulated inhibitory phosphorylation of the KCC2 cotransporter maintains the depolarizing action of GABA in immature neurons. Sci Signal 8:ra65.

Gackiere F, Vinay L (2015) Contribution of the potassium-chloride cotransporter KCC2 to the strength of inhibition in the neonatal rodent spinal cord in vitro. J Neurosci 35:5307–5316.

Gagnon M, Bergeron MJ, Lavertu G, Castonguay A, Tripathy S, Bonin RP,et al. Chloride extrusion enhancers as novel therapeutics for neurological diseases. Nat Med 2013; 19: 1524–8.

Gomez-Pinilla F, Huie JR, Ying Z, Ferguson AR, Crown ED, Baumbauer KM,et al. BDNF and learning: Evidence that instrumental training promotes learning within the spinal cord by up-regulating BDNF expression. Neuroscience 2007; 148: 893–906.

Grey MJ, Klinge K, Crone C, Lorentzen J, Biering-Sorensen F, Ravnborg M,et al. Post-activation depression of soleus stretch reflexes in healthy and spastic humans. Exp Brain Res 2008; 185: 189–97.

Harkema SJ. Plasticity of interneuronal networks of the functionally isolated human spinal cord. Brain Res Rev 2008; 57: 255–64.

Hasbargen T, Ahmed MM, Miranpuri G, Li L, Kahle KT, Resnick D, Sun D (2010) Role of NKCC1 and KCC2 in the development of chronic neuropathic pain following spinal cord injury. AnnNYAcadSci 1198:168–172.

Holtz KA, Lipson R, Noonan VK, Kwon BK, Mills PB. Prevalence and Effect of Problematic Spasticity After Traumatic Spinal Cord Injury. Arch Phys Med Rehabil 2017; 98: 1132–8.

Houle JD, Morris K, Skinner RD, Garcia-Rill E, Peterson CA. Effects of fetal spinal cord tissue transplants and cycling exercise on the soleus muscle in spinalized rats. Muscle Nerve 1999; 22: 846–56.

Huang YJ, Lee KH, Grau JW (2017) Complete spinal cord injury (SCI) transforms how brain derived neurotrophic factor (BDNF) affects nociceptive sensitization. Exp Neurol 288:38–50.

Hübner CA, Stein V, Hermans-Borgmeyer I, Meyer T, Ballanyi K, Jentsch TJ. Disruption of KCC2 reveals an essential role of K-Cl cotransport already in early synaptic inhibition. Neuron 2001; 30: 515–24.

Huie JR, Garraway SM, Baumbauer KM, Hoy KC, Jr., Beas BS, Montgomery KS, Bizon JL, Grau JW (2012) Brain-derived neurotrophic factor promotes adaptive plasticity within the spinal cord and mediates the beneficial effects of controllable stimulation. Neuroscience 200:74–90.

Hutchinson KJ, Gomez-Pinilla F, Crowe MJ, Ying Z, Basso DM. Three exercise paradigms differentially improve sensory recovery after spinal cord contusion in rats. Brain 2004; 127(Pt 6): 1403–14.

Jean-Xavier C, Pflieger JF, Liabeuf S, Vinay L. Inhibitory postsynaptic potentials in lumbar motoneurons remain depolarizing after neonatal spinal cord transection in the rat. J Neurophysiol 2006; 96: 2274–81.

Jolivalt CG, Lee CA, Ramos KM, Calcutt NA. Allodynia and hyperalgesia in diabetic rats are mediated by GABA and depletion of spinal potassium-chloride co-transporters. Pain 2008; 140: 48–57.

Kahle KT, Khanna A, Clapham DE, Woolf CJ. Therapeutic restoration of spinal inhibition via druggable enhancement of potassium-chloride cotransporter KCC2-mediated chloride extrusion in peripheral neuropathic pain. JAMA Neurol 2014; 71: 640–5.

Kahle KT, Rinehart J, Lifton RP. Phosphoregulation of the Na-K-2Cl and K-Cl cotransporters by the WNK kinases. Biochim Biophys Acta 2010; 1802: 1150–8.

Kahle KT et al. (2016) Inhibition of the kinase WNK1/HSN2 ameliorates neuropathic pain by restoring GABA inhibition. Sci Signal 9:ra32.

Kaila K, Price TJ, Payne JA, Puskarjov M, Voipio J. Cation-chloride cotransporters in neuronal development, plasticity and disease. Nat Rev Neurosci 2014a; 15: 637–54.

Kaila K, Ruusuvuori E, Seja P, Voipio J, Puskarjov M. GABA actions and ionic plasticity in epilepsy. Curr Opin Neurobiol 2014b; 26: 34–41.

Katz R. Presynaptic inhibition in humans: a comparison between normal and spastic patients. J Physiol (Paris) 1999; 93: 379–85.

Khirug S, Ahmad F, Puskarjov M, Afzalov R, Kaila K, Blaesse P. A single seizure episode leads to rapid functional activation of KCC2 in the neonatal rat hippocampus. J Neurosci 2010; 30: 12028–35.

Lee HH, Walker JA, Williams JR, Goodier RJ, Payne JA, Moss SJ. Direct protein kinase C-dependent phosphorylation regulates the cell surface stability and activity of the potassium chloride cotransporter KCC2. J Biol Chem 2007; 282: 29777–84.

Lee-Kubli CA, Calcutt NA. Altered rate-dependent depression of the spinal H-reflex as an indicator of spinal disinhibition in models of neuropathic pain. Pain 2014; 155: 250–60.

Leech KA, Hornby TG. High-Intensity Locomotor Exercise Increases Brain-Derived Neurotrophic Factor in Individuals with Incomplete Spinal Cord Injury. J Neurotrauma 2017; 34: 1240–8.

Lu B. BDNF and activity-dependent synaptic modulation. Learn Mem 2003; 10: 86–98.

Lu Y, Zheng J, Xiong L, Zimmermann M, Yang J. Spinal cord injury-induced attenuation of GABAergic inhibition in spinal dorsal horn circuits is associated with down-regulation of the chloride transporter KCC2 in rat. J Physiol 2008; 586: 5701–15.

Ludwig A, Uvarov P, Soni S, Thomas-Crusells J, Airaksinen MS, Rivera C. Early growth response 4 mediates BDNF induction of potassium chloride cotransporter 2 transcription. J Neurosci 2011; 31: 644–9.

Maynard FM, Karunas RS, Waring WP, 3rd. Epidemiology of spasticity following traumatic spinal cord injury. Arch Phys Med Rehab 1990; 71: 566–9.

Mazzocchio R, Rossi A. Involvement of spinal recurrent inhibition in spasticity. Further insight into the regulation of Renshaw cell activity. Brain 1997; 120 (Pt 6): 991–1003.

Miletic G, Miletic V. Loose ligation of the sciatic nerve is associated with TrkB receptor-dependent decreases in KCC2 protein levels in the ipsilateral spinal dorsal horn. Pain 2008; 137: 532–9.

Mòdol L, Mancuso R, Ale A, Francos-Quijorna I, Navarro X. Differential effects on KCC2 expression and spasticity of ALS and traumatic injuries to motoneurons. Front Cell Neurosci 2014b; 8: 7.

Payne JA, Rivera C, Voipio J, Kaila K. Cation-chloride co-transporters in neuronal communication, development and trauma. Trends Neurosci 2003; 26: 199–206.

Petropoulou KB, Panourias IG, Rapidi CA, Sakas DE. The importance of neurorehabilitation to the outcome of neuromodulation in spasticity. Acta Neurochir Suppl 2007; 97(Pt 1): 243–50.

Plantier V, Brocard F. [Calpain as a new therapeutic target for treating spasticity after a spinal cord injury]. Med Sci (Paris) 2017; 33(6-7): 629–36.

Puskarjov M, Ahmad F, Kaila K, Blaesse P. Activity-dependent cleavage of the K-Cl cotransporter KCC2 mediated by calcium-activated protease calpain. J Neurosci 2012; 32: 11356–64.

Puskarjov M, Ahmad F, Khirug S, Sivakumaran S, Kaila K, Blaesse P. BDNF is required for seizure-induced but not developmental up-regulation of KCC2 in the neonatal hippocampus. Neuropharmacology 2015; 88: 103–9.

Rivera C, Li H, Thomas-Crusells J, Lahtinen H, Viitanen T, Nanobashvili A,et al. BDNF-induced TrkB activation down-regulates the K+-Cl-cotransporter KCC2 and impairs neuronal Cl-extrusion. J Cell Biol 2002; 159: 747–52.

Rivera C, Voipio J, Payne JA, Ruusuvuori E, Lahtinen H, Lamsa K,et al. The K+/Cl-cotransporter KCC2 renders GABA hyperpolarizing during neuronal maturation. Nature 1999; 397: 251–5.

Rivera C, Voipio J, Thomas-Crusells J, Li H, Emri Z, Sipila S,et al. Mechanism of activity-dependent downregulation of the neuron-specific K-Cl cotransporter KCC2. J Neurosci 2004; 24: 4683–91.

Skold C, Levi R, Seiger A. Spasticity after traumatic spinal cord injury: nature, severity, and location. Arch Phys Med Rehabil 1999; 80: 1548–57.

Skup M, Ziemlinska E, Gajewska-Wozniak O, Platek R, Maciejewska A, Czarkowska-Bauch J. The impact of training and neurotrophins on functional recovery after complete spinal cord transection: cellular and molecular mechanisms contributing to motor improvement. Acta Neurobiol Exp (Wars) 2014; 74: 121–41.

Tan T, Watts SW, Davis RP. Drug Delivery: Enabling Technology for Drug Discovery and Development. iPRECIO Micro Infusion Pump: Programmable, Refillable, and Implantable. Front Pharmacol 2011; 2: 44.

Tashiro S, Shinozaki M, Mukaino M, Renault-Mihara F, Toyama Y, Liu M,et al. BDNF Induced by Treadmill Training Contributes to the Suppression of Spasticity and Allodynia After Spinal Cord Injury via Upregulation of KCC2. Neurorehabil Neural Repair 2015; 29: 677–89.

Thompson FJ, Reier PJ, Lucas CC, Parmer R. Altered patterns of reflex excitability subsequent to contusion injury of the rat spinal cord. J Neurophysiol 1992; 68: 1473–86.

Weishaupt N, Blesch A, Fouad K. BDNF: the career of a multifaceted neurotrophin in spinal cord injury. Exp Neurol 2012; 238: 254–64.

Yates C, Charlesworth A, Allen SR, Reese NB, Skinner RD, Garcia-Rill E. The onset of hyperreflexia in the rat following complete spinal cord transection. Spinal Cord 2008; 46: 798–803.

Ying Z, Roy RR, Zhong H, Zdunowski S, Edgerton VR, Gomez-Pinilla F. BDNF-exercise interactions in the recovery of symmetrical stepping after a cervical hemisection in rats. Neuroscience 2008; 155: 1070–8.

Zadran S, Jourdi H, Rostamiani K, Qin Q, Bi X, Baudry M Brain-derived neurotrophic factor and epidermal growth factor activate neuronal m-calpain via mitogen-activated protein kinase-dependent phosphorylation. J Neurosci 2010; 30: 1086–95.

Zhou HY, Chen SR, Byun HS, Chen H, Li L, Han HD,et al. N-methyl-D-aspartate receptor-and calpain-mediated proteolytic cleavage of K+-Cl-cotransporter-2 impairs spinal chloride homeostasis in neuropathic pain. J Biol Chem 2012; 287: 33853–64.

